# Insular hemorrhagic stroke in mice: a model of neurocardiac dysfunction

**DOI:** 10.64898/2026.08.03.741256

**Authors:** A. C. Ventris-Godoy, H. Abramo, L. Rodrigues-Ribeiro, A. C. Rocha-Viana, G. Pires, R. A. S. Santos, C. Rocha-Resende, M. A. P. Fontes

**Affiliations:** ¹Department of Physiology and Biophysics, Federal University of Minas Gerais (UFMG), Belo Horizonte/ MG; ²Department of Biochemistry and Immunology, Federal University of Minas Gerais (UFMG), Belo Horizonte/ MG

**Author notes:** Corresponding author: M.A.P. Fontes, Ph.D. Hypertension Laboratory, INCT NanoBiofar, Department of Physiology and Biophysics ICB, Federal University of Minas Gerais. Belo Horizonte, Brazil, MG 31270 901. Co-mentorship; these authors jointly supervised this work (C. Rocha-Resende, M. A. P. Fontes).

**Keywords:** Insula, Cortex, Stroke, Autonomic, Rilmenidine, Cardiovascular, Mice

## Abstract

**Background:** Insular damage leads to marked cardiovascular alterations and the mechanisms need to be understood. Mouse models provide unique opportunities to gain insights into pathophysiological mechanisms. Here, we evaluated the effects of rilmenidine, a centrally acting antihypertensive drug, on the cardiac functional parameters and cardiac inflammatory cell infiltration in a newly developed mice model of insular hemorrhagic stroke.

**Methods:** C57BL/6J mice were instrumented for injection of blood or vehicle into the insular cortex (IC). Immediately after IC stroke induction, separate groups received intraperitoneal treatment with vehicle (0.9% NaCl, 0.1 mL/100 g) or rilmenidine (10 μg/kg) for three days. Electrocardiogram recording,cardiac catecholamine levels and myocardial accumulation of immune cells were evaluated.

**Results:** Mice subjected to hemorrhagic stroke exhibited higher baseline heart rate (HR) (control: 296 ± 33 bpm vs. stroke: 349 ± 38 bpm; *P* < 0.01) and prolonged QTc interval (control: 89 ± 11 ms vs. stroke: 100 ± 7 ms; *P* < 0.01). Stroke also increased cardiac norepinephrine levels (control: 9 ± 4 ng/mg vs. stroke: 25 ± 14 ng/mg; *P* < 0.05), as well as the number of myocardial CD68+ macrophages (control: 7 ± 4 vs. stroke: 16 ± 6 cells/field; *P* < 0.0001) and Ly6G+ neutrophils (control: 0.5 ± 0.7 vs. stroke: 1.5 ± 1 cells/field; *P* < 0.001). Rilmenidine treatment markedly prevented all major stroke-induced myocardial functional and inflammatory changes

**Conclusions:** Insular hemorrhagic stroke in mice induces centrally mediated cardiac noradrenergic hyperactivation accompanied by myocardial accumulation of immune cells. These findings support the relevance of this murine model for investigating mechanisms associated with insular stroke.

## Introduction

Stroke represents one of the leading causes of global morbidity and mortality, with hemorrhagic stroke standing out due to its severity and high lethality (Wang, 2010; Campbell et al., 2019; Feigin et al., 2022). The location of the brain lesion is a determining factor for prognosis (Wang, 2010; Lai et al., 2018; Campbell et al., 2019). In particular, the insular cortex (IC) plays a central role in autonomic regulation (Christensen, 2005; Rincon et al., 2008; Oppenheimer and Cechetto, 2016). Hemorrhagic stroke in the IC, especially in the right hemisphere, are associated with severe cardiovascular dysfunctions, including arrhythmias, ventricular dysfunction, and a significant increase in mortality, particularly in the first months post-stroke (Oppenheimer et al., 1991a; Wittling et al., 1998; Watkins, 2001; Meyer et al., 2004; Christensen, 2005; Rincon et al., 2008; Seifert et al., 2015; Oppenheimer and Cechetto, 2016; Laredo et al., 2018; Marins et al., 2020; Sanchez-Larsen et al., 2021; Sanchez-Larsen et al., 2024).

Previous studies by our group have demonstrated a rostrocaudal functional topography in the IC for autonomic control, with a relatively distinct sympathoexcitatory area (Marins et al., 2016). Furthermore, we confirmed hemispheric asymmetry, observing a significantly higher incidence of arrhythmias after stroke in the right insula of rats compared to the left insula (Marins et al., 2020; Dos Santos Machado et al., 2026). Corroborating these experimental findings, recent clinical studies also associate insular lesions with cardiac tissue damage, impaired left ventricular ejection fraction, and an increased risk of adverse cardiac events (Winder et al., 2023).

The investigation of hemorrhagic stroke pathophysiological mechanisms and the development of effective therapeutic strategies depend on robust experimental models. Traditionally, ischemic stroke models are widely used, but they often result in extensive and non-specific lesions, frequently reaching subcortical regions essential for cardiovascular control, which compromises the effectiveness of investigations focused on the IC (Sommer, 2017; Bai et al., 2020). In contrast, the use of a hemorrhagic stroke model restricted to the IC allows for the reproduction of clinically observed cardiovascular autonomic changes, with focal and controlled lesions (Marins et al., 2020; Dos Santos Machado et al., 2026). The use of C57BL/6J mice, a genetically well-characterized strain and basis for numerous transgenic and knockout models, represents a valuable platform for investigations associated with post-stroke central autonomic dysfunction resulting from insular damage (Kleinschnitz et al., 2015).

In this context, rilmenidine, a second-generation centrally acting antihypertensive, presents itself as a potential therapeutic agent. Its pharmacological action is mediated by the selective activation of I1 imidazoline receptors, predominantly located in the rostral ventrolateral medulla (RVLM) – a region essential for the tonic control of sympathetic activity (Ehrhardt, 1985; LAUBIE et al., 1985; Bricca et al., 1988; Bousquet et al., 1989). The activation of I1 RVLM receptors by rilmenidine promotes a sympatholytic effect, resulting in vasodilation, reduced peripheral resistance, sustained decrease in blood pressure, accompanied by bradycardia, in addition to reducing circulating catecholamine levels (Bousquet and Guertzenstein, 1973; Bousquet et al., 1981, 1984; Bricca et al., 1988; Valet et al., 1988; Feldman et al., 1990; Esler et al., 2004). Rilmenidine selectivity for I1 receptors minimizes the typical adverse effects of α2-adrenergic agonists, such as sedation and dry mouth, representing an important promising therapeutic option (Bousquet et al., 2000; Mahmoudi et al., 2018).

The present study characterizes an experimental hemorrhagic stroke model restricted to the IC in C57BL/6J mice, using a focal and controlled volume of autologous blood. The objective is to evaluate the cardiovascular alterations in this model and the therapeutic potential of rilmenidine. The hypothesis of this study is that hemorrhagic stroke in the insula will induce cardiac sympathetic hyperactivity and that rilmenidine administration will attenuate these cardiovascular dysfunctions.

## Materials and Methods

### Animals

Male C57BL/6J mice (20-25g, 10-12 weeks old) were obtained from the Central Animal Facility of the Federal University of Minas Gerais (UFMG). Animals were housed in polypropylene cages with wood shavings and maintained under controlled temperature (22-24°C) on a 12/12-h light-dark cycle. Standard granulated chow and water were provided *ad libitum*. All experimental procedures were performed in accordance with protocol n°233/2023, approved by the Institutional Animal Care and Use Committee at UFMG, and followed the National Institutes of Health (NIH) Guide for the Care and Use of Laboratory Animals.

### Drugs and solutions

For surgical procedures, mice were anesthetized with an intraperitoneal (i.p.) injection of ketamine (80 mg/kg) and xylazine (7 mg/kg). Following stereotaxic, a single dose of veterinary pentabiotic (40,000 IU/kg, Fort Dodge®) was administered intramuscularly (i.m.), and flunixin meglumine (2.5 mg/kg, Banamine®) was given subcutaneously (s.c.) for post-operative analgesia. Asepsis was maintained using 2% chlorhexidine (Riohex®) and 70% alcohol. Rilmenidine (Sigma-Aldrich, USA) was administered at a dose of 10 µg/kg. Control subjects received sterile saline (0.9% NaCl; 0.1 ml/100g) i.p., and a 100 nL microinjection of saline was also delivered directly into the right IC.

### Hemorrhagic stroke induction

Mice were anesthetized with a ketamine/xylazine mixture (87/13 mg/kg, i.p.) and secured in a small-animal stereotaxic apparatus (Stoelting, USA). Following cranial depilation and local asepsis, a 0.5 cm incision was made to expose the skull, and the periosteum was removed. To induce intracerebral hemorrhage (ICH), as previously described (Marins et al., 2020; Dos Santos Machado et al., 2026), autologous blood was microinjected into the right IC. Blood was collected from a tail incision after local vasodilation induced by a thermal gel pack (Termogel®). The collected blood was transferred to a 30G needle connected to a 1 µL nanoinjector (Hamilton, USA) via Silastic® tubing. A volume of 100 nL was injected at the following coordinates (Paxinos and Franklin, 2004): AP -0.10 mm from bregma; ML +3.7 mm from midline; and DV 4.5 mm from the skull surface. Control mice for stroke received an equal volume (100 nL) of sterile saline into the insular cortex. Following suturing, subjects received their respective first day treatments (intraperitoneal rilmenidine or saline). Post-operative care included prophylactic pentabiotic and flunixin meglumine, with recovery monitored in a heated environment under an infrared lamp (Philips®). Treatments were continued on the second and third days post-induction.

### Electrocardiography (ECG)

Under ketamine/xylazine anesthesia, copper wire electrodes were inserted subcutaneously into the limbs and connected to a PowerLab 4/20 acquisition system (ADInstruments). Data was captured and analyzed using LabChart 8.1.31 software. Cardiac electrical activity was recorded for 5 minutes per subject in the ventral decubitus position. Body temperature was maintained at 37°C using a heating platform and monitored via a rectal probe. Heart rate and electrocardiographic tracings were subsequently analyzed.

### Immunofluorescence

Immunofluorescence was performed as previously described Rocha-Resende et al.(2019). Hearts were fixed in 4% PFA, embedded in OCT, and sectioned (10 µm) using a cryostat. Sections were incubated in blocking buffer (20% Fetal Bovine Serum and 2% Roche Blocking Reagent, Roche; Catalog #11096176001) for 2 hours at room temperature, followed by overnight incubation at 4°C with primary antibodies: Rat anti-mouse CD68 FA-11 (1:300, BioRad, Catalog #MAC1957, RRID:AB_322219) for macrophages and Purified Rat Anti-Mouse Ly-6G 1A8 (1:200, BD Biosciences Cat# 551459, RRID:AB_394206) for neutrophils. Secondary antibodies (1:200, Alexa Fluor 488 goat anti-rat, Invitrogen, Catalog #A11006, RRID:AB_2534074, or Alexa Fluor 555 goat anti-rat,, Invitrogen, Catalog #A21434, RRID:AB_2535855) were applied for 2 hours at room temperature. Nuclei were counterstained with DAPI. Images were acquired using an Zeiss APOTOME II microscope at 20x magnification. For each subject, 10 images were analyzed across two distinct sections using ZEN 3.6 (blue edition) software. Cell quantification was expressed as the number of CD68^+^DAPI^+^ or Ly-6G^+^DAPI^+^ cells per field.

### Cardiac catecholamine quantification

Hearts were harvested following cervical dislocation, cleaned in PBS, snap-frozen in liquid nitrogen, and stored at −80°C. Tissue was resuspended in a deproteinization solution (0.2 M perchloric acid, 3 mM cysteine), sonicated on ice, and centrifuged (15,000 × g, 15 min, 4°C). The supernatant was analyzed via high-performance liquid chromatography with fluorescence detection (HPLC-FD, Shimadzu) using a C18 reverse-phase column (250 × 4.6 mm, 5 μm; Phenomenex). The mobile phase (12 mM sodium acetate, 0.26 mM disodium EDTA, pH 3.5) was delivered isocratically at 0.5 mL/min. Catecholamines were detected at λₑₓ = 279 nm and λₑₘ = 320 nm.

### Experimental design

The complete experimental design is presented in Figure 1. Mice were transported to the experimental room in their home cages and allowed to acclimatize for approximately 60 minutes. The experimental room was climate-controlled (± 25°C). All animals initially underwent a surgical procedure to induce intracerebral hemorrhage targeted at the right intermediate IC via microinjection of autologous blood; volume control subjects received an equal volume of saline into the insular cortex (0.9% NaCl). Animals were subjecte to three days of treatment of either intraperitoneal rilmenidine (10 µg/kg) or saline vehicle (0.9% NaCl, 0,1 ml/100 g): the first immediately after surgery, followed by subsequent doses on days 2 and 3 post-stroke or stroke control. After a 24- hour period to adapt to the experimental room conditions (e.g., sound and lighting), animals were anesthetized and underwent a 5-minute ECG recording. Following the recording, subjects were euthanized by cervical dislocation. Hearts were immediately harvested for immunohistochemical and biochemical analyses, and brains were collected for histological verification of the injection sites.

**Figure 1.**
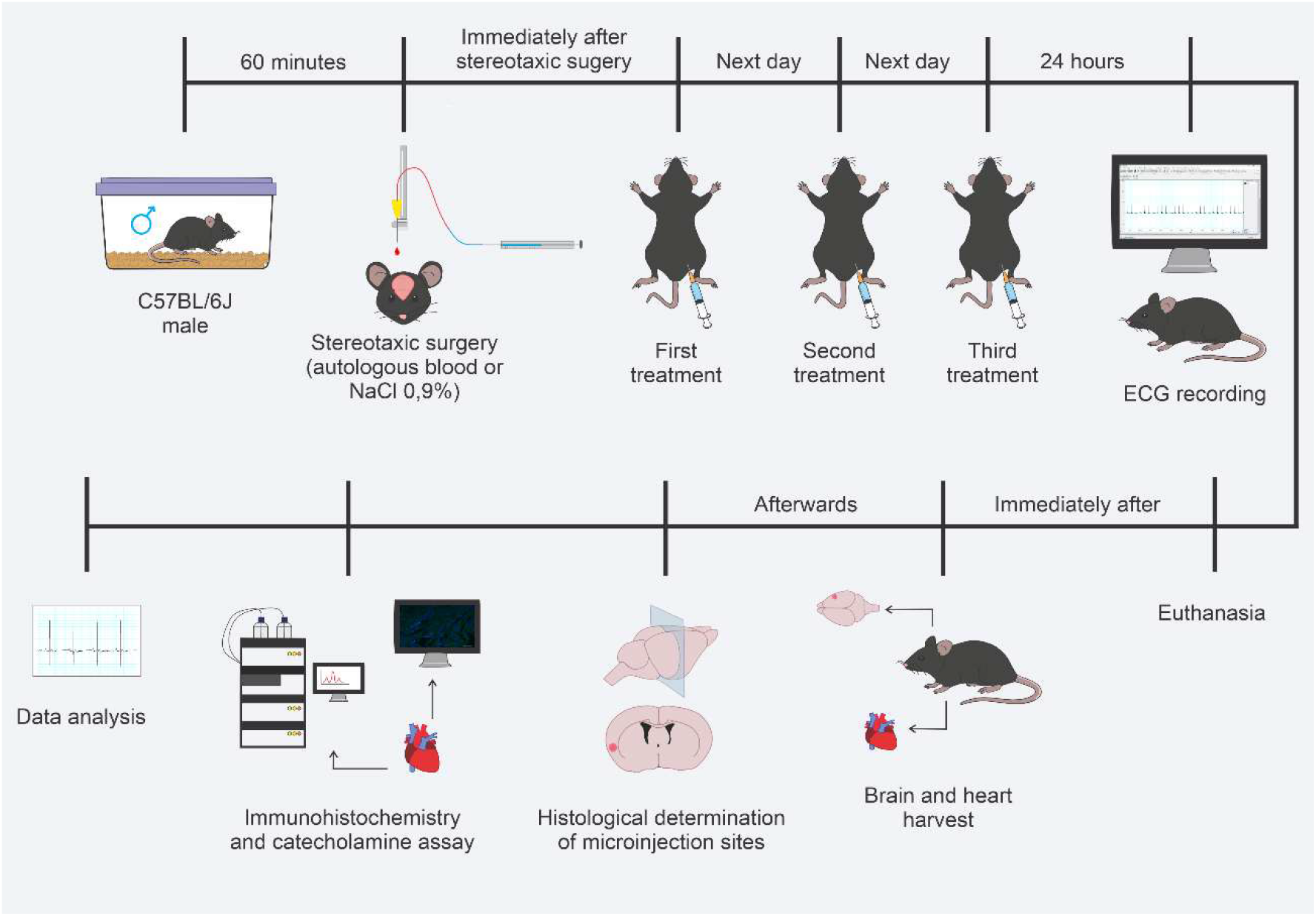
Experimental protocol. Schematic representation of the experimental procedures: 1) groups subjected to stroke induction (100 nL of autologous blood injected into the insular cortex) or central volume control (100 nL of NaCl 0,9 % injected into the insular cortex); 2) post-stroke treatments (rilmenidine, 10 µg/kg i.p. or saline vehicle 0.9% NaCl, 0,1 ml/100g, i.p.); 3) ECG recordings, insular cortex injection site confirmation (histological analysis), tissue harvesting, and subsequent overall data analyses.

### Histological determination of microinjection sites

Following experimental procedures, brains were harvested, fixed in 10% buffered formalin for 24 hours, and cryoprotected in 20% sucrose for 24-48 hours. Frontal sections (50 μm) were obtained using a cryostat (LEICA CM 1850®). Microinjection sites were verified by serial section analysis and identified according to the mouse brain atlas (Paxinos and Franklin, 2004).

### Data and statistical analysis

Data were analyzed using GraphPad Prism 8.0.1 and are presented as mean ± standard error of the mean (SEM). Inter-group comparisons were performed using one-way ANOVA followed by Tukey’s post-hoc test. Statistical significance was set at p < 0.05.

## Results

Histological analysis of the microinjection sites confirmed the precise localization within the IC. Figure 2 illustrates the topographical mapping along the anteroposterior axis of the IC. Based on our previous studies demonstrating that stroke in the right intermediate IC induces greater cardiac dysfunction, microinjections were specifically targeted at the intermediate axis of the right IC.

**Figure 2.**
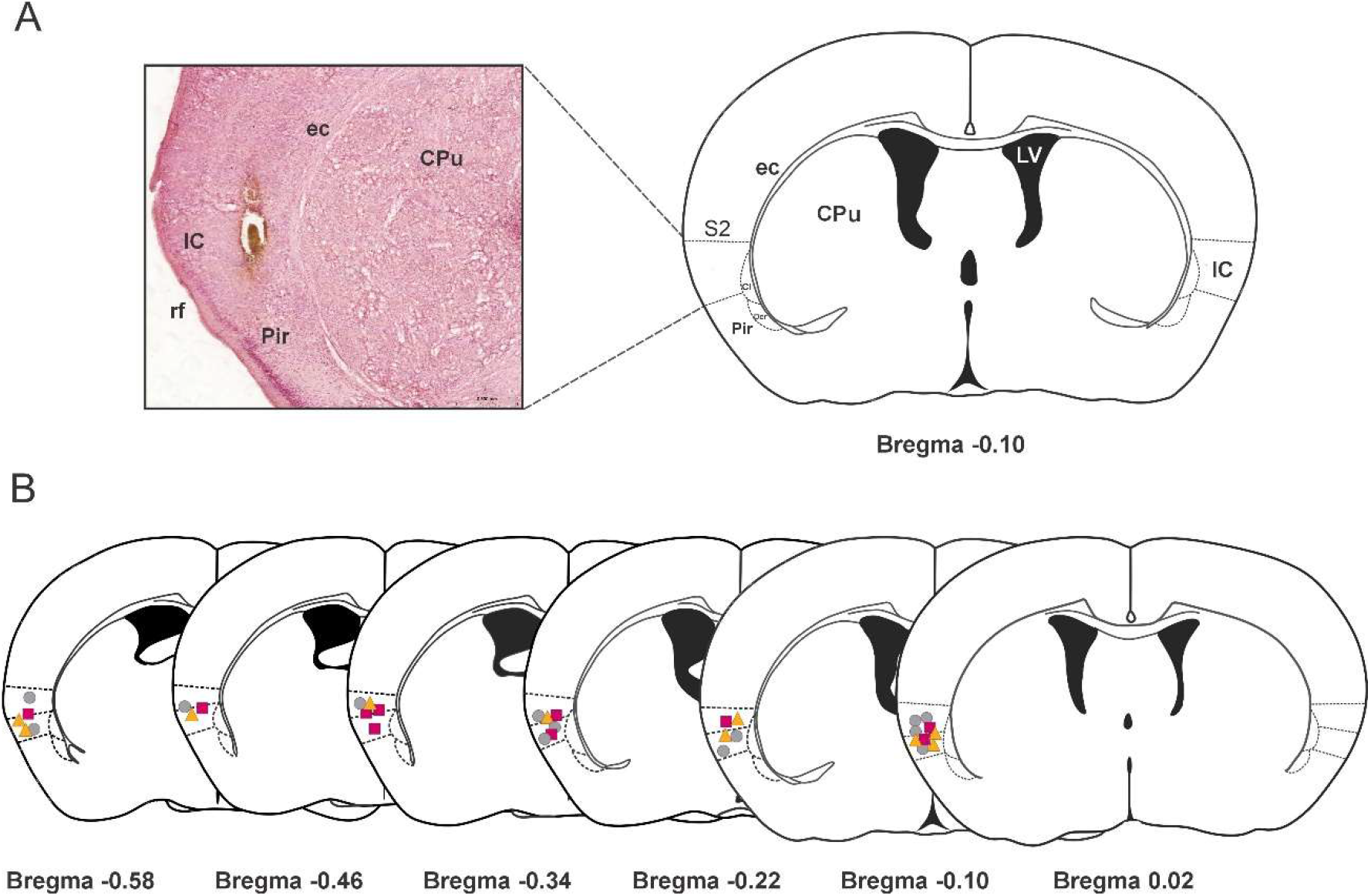
Topographical mapping mouse brain depicting insular injection sites. **(A)** Representative photomicrograph of the insular cortex (IC) (frontal view) at the stereotaxic coordinate: AP -0.10 mm from bregma (drawing based on Paxinos and Franklin, 2004) in a mouse following stroke induction. The injection site is shown at 2.4x magnification using SlideViewer software (version 2.9). **(B)** Diagrammatic representation indicating the distribution of injection sites for saline (gray circles), stroke (pink squares), and stroke-rilmenidine (yellow triangles) within the IC. S2, secondary somatosensory cortex; ec, external capsule; Cl, claustrum; DEn, dorsal endopiriform nucleus; Pir, piriform cortex; rf, rhinal fissure; IC, insular cortex; CPu, caudate-putamen (striatum); LV, lateral ventricle.

Microinjection of autologous blood into the IC of C57BL/6J mice induced sustained tachycardia (control: 296 ± 33 bpm vs. stroke: 349 ± 38 bpm; P < 0.01). Treatment with rilmenidine (10 µg/kg) prevented the development of tachycardia (stroke- rilmenidine: 310 ± 22 bpm; P < 0.05 compared to stroke) (Figure 3A). ECG analysis revealed a significant increase in the QTc interval in mice following insular stroke (control: 89 ± 11 ms vs. stroke: 100 ± 7 ms; P < 0.01). Rilmenidine treatment did not significantly attenuate this QTc prolongation, with the treated group showing no statistical difference from either the stroke or control groups (stroke-rilmenidine: 96 ± 5 ms; P > 0.05) (Figure 3B). Representative ECG tracings for each group are shown in Figure 3C.

**Figure 3.**
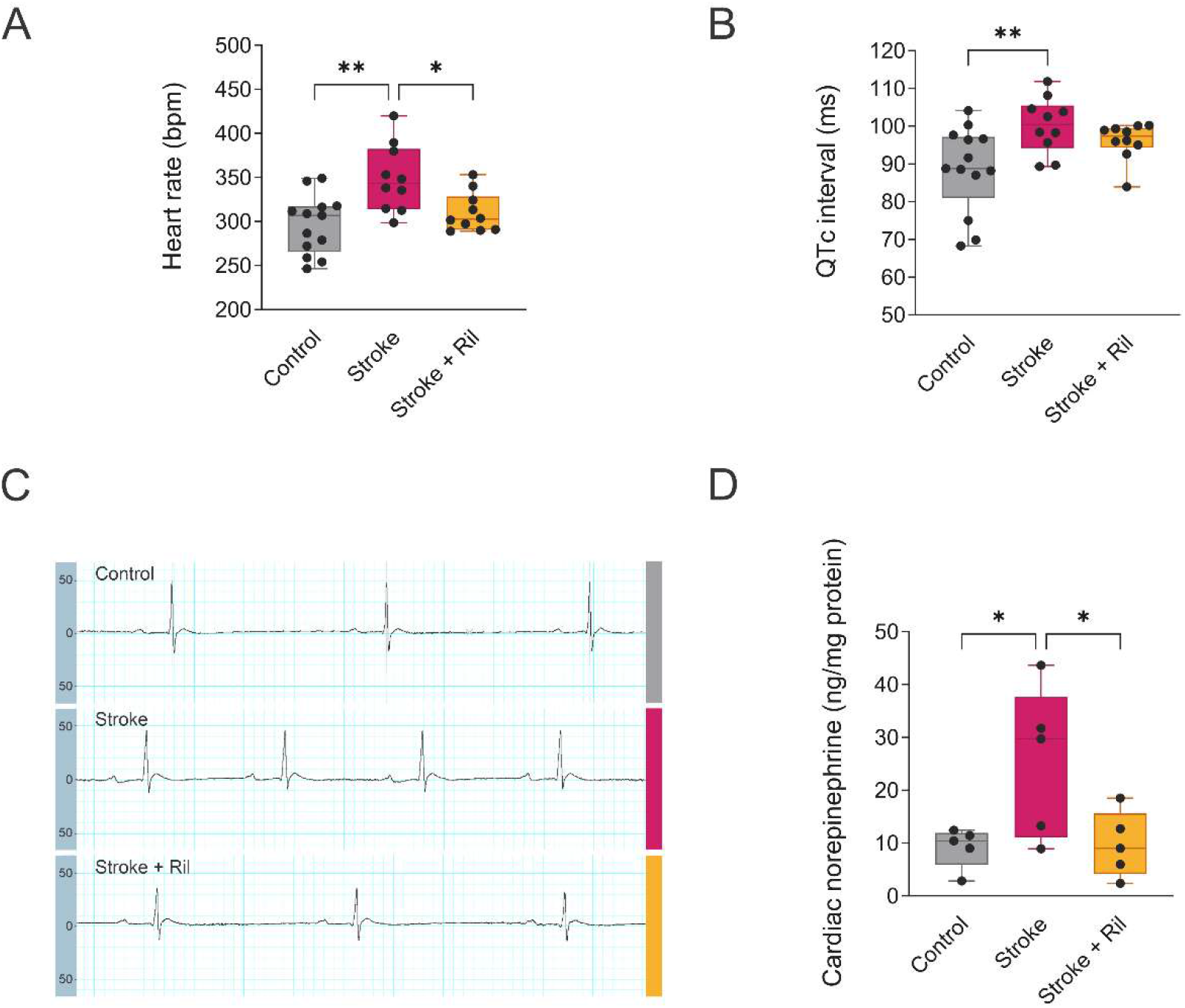
Cardiovascular and biochemical effects of experimental stroke in the insular cortex (IC). Analysis of control animals (gray bars; 100 nL saline in the IC + saline i.p., n = 13), stroke animals (pink bars; 100 nL autologous blood in the IC + saline i.p., n = 10), or stroke-rilmenidine animals (yellow bars; 100 nL autologous blood in the IC + 10 µg/kg rilmenidine i.p., n = 10). **(A)** Heart rate analysis in beats per minute (bpm). **(B)** QTc interval analysis in milliseconds (ms). **(C)** Representative ECG tracings. **(D)** Cardiac norepinephrine quantification in ng/mg of protein (n = 5 per group). Black circles represent individual data points. The values are represented as the means ± standard error of the mean (SEM). Statistical analysis was performed using one-way ANOVA followed by Tukey’s multiple comparisons test. *p < 0.05 and **p < 0.01 indicate significant differences between the indicated groups.

To assess peripheral sympathetic hyperactivation induced by the stroke, cardiac norepinephrine concentrations were quantified. Norepinephrine levels were significantly elevated in the hearts of stroke mice (control: 9 ± 4 ng/mg; stroke: 25 ± 14 ng/mg; P < 0.05), reflecting sympathetic hyperactivity. Rilmenidine treatment (10 µg/kg) reversed this increase, restoring norepinephrine levels to values similar to those of the control group (stroke-rilmenidine: 8 ± 6 ng/mg; P < 0.05 compared to stroke) (Figure 3D).

Given the stroke-induced tachycardia and its prevention by rilmenidine, and considering that peripheral sympathetic hyperactivation can induce myocardial damage, we investigated the accumulation of CD68^+^ macrophages in cardiac tissue. Results revealed a mild, but significant, increase in the number of CD68^+^ macrophages in mice subjected to stroke compared to the control group (control: 7 ± 4 vs. stroke: 16 ± 6 cells/field; P < 0.0001). Rilmenidine treatment prevented this increase (stroke- rilmenidine: 10 ± 5 cells/field; P < 0.0001, compared to stroke) (Figure 4A-B). CD68^+^ macrophages in stroke group were founddiffusely throughout the parenchyma and, in certain regions, aggregated in clusters, indicating an increased number of infiltrating CD68^+^ macrophages (Figure 4A-B).

**Figure 4.**
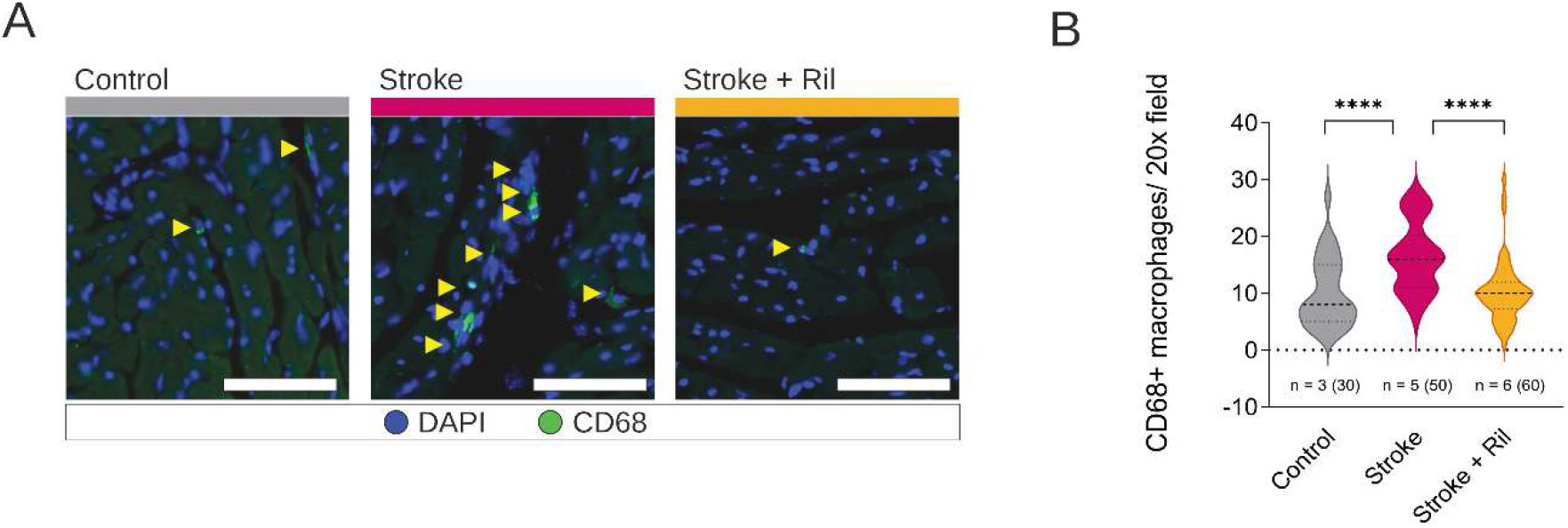
Cardiac CD68^+^ macrophage infiltration following insular stroke. **(A)** Representative immunofluorescence images of CD68^+^ macrophages in the heart. Green, CD68; blue, DAPI. Scale bar = 25 μm. **(B**) Bar graph showing the quantification of CD68^+^ macrophages per 20x field (control, gray bars, n = 3 mice/30 fields; stroke, pink bars, n = 5 mice/50 fields; stroke + rilmenidine, yellow bars, n = 6 mice/60 fields). Bars represent the mean, and error bars indicate the standard error of the mean (± SEM). Statistical analysis was performed using one-way ANOVA followed by Tukey’s multiple comparisons test. ****p < 0.0001 indicate significant differences between the indicated groups.

Furthermore, we investigated the presence of Ly6G^+^ neutrophils, as their elevation is also associated with the myocardial inflammatory response. As shown in Figure 5A-B, there was an increase in the number of Ly6G^+^ cells in stroke mice compared to the control group (control: 0.5 ± 0.7 vs. stroke: 1.5 ± 1 cells/field; P < 0.001). Similar to the observations for CD68^+^, rilmenidine treatment prevented neutrophil infiltration (stroke-rilmenidine: 0.7 ± 0.9 cells/field; P < 0.001, compared to stroke), as evidenced in Figure 5A–B.

**Figure 5.**
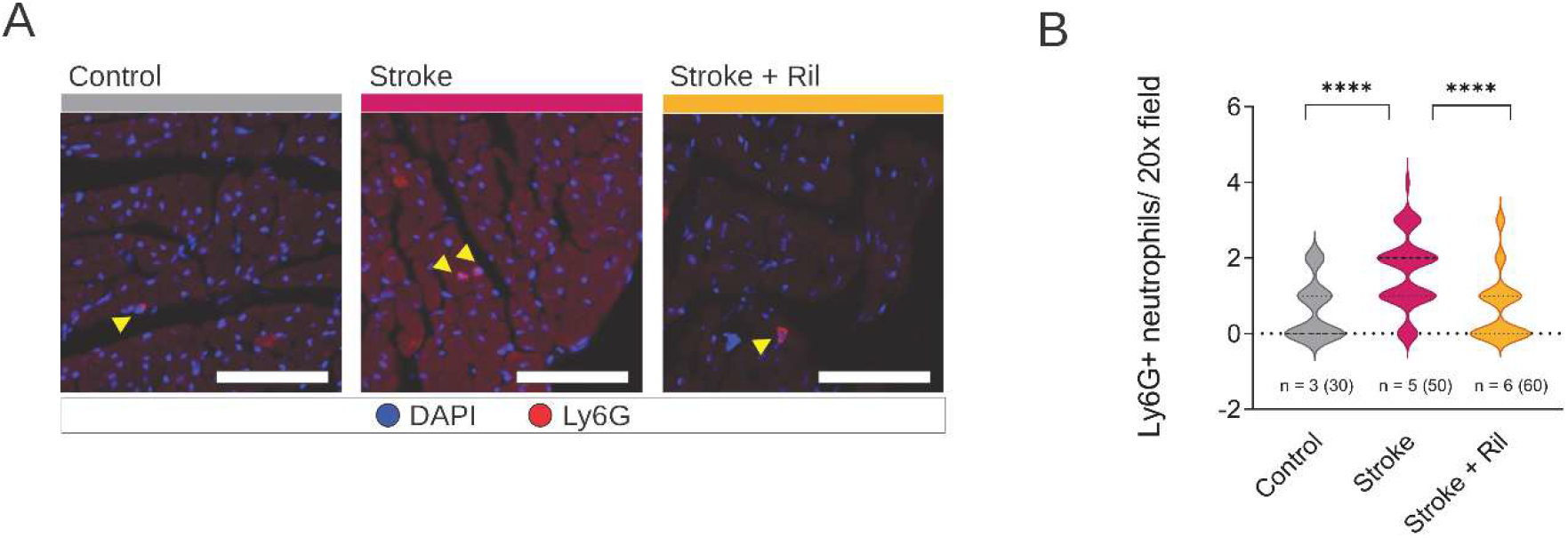
Cardiac Ly6G^+^ neutrophil infiltration following insular stroke. **(A)** Representative immunofluorescence images of Ly6G^+^ neutrophils in the heart. Red, Ly6G; blue, DAPI. Scale bar = 25 μm. **(B**) Bar graph showing the quantification of Ly6G^+^ neutrophils per 20x field (control, gray bars, n = 3 mice/30 fields; stroke, pink bars, n = 5 mice/50 fields; stroke + ril, yellow bars, n = 6 mice/60 fields). Bars represent the mean, and error bars indicate the standard error of the mean (± SEM). Statistical analysis was performed using one-way ANOVA followed by Tukey’s multiple comparisons test. ****p < 0.001 indicates significant differences between the indicated groups.

## Discussion

Stroke involving the insular cortex (IC) is strongly associated with cardiovascular complications in patients, yet experimental models that faithfully reproduce this neurocardiac interaction remain limited. In the present study, we established a novel murine model of insular hemorrhagic stroke that recapitulates key features of stroke-induced autonomic and cardiac dysfunction. Animals exhibited sustained tachycardia, QTc interval prolongation, and increased cardiac norepinephrine content, providing direct evidence of enhanced sympathetic activity directed to the heart. These autonomic alterations were accompanied by increased accumulation of macrophages and neutrophils in the heart, suggesting that sympathetic hyperactivity is associated not only with electrophysiological disturbances but also with the development of cardiac inflammation. Importantly, tachycardia and the increase in immune cells in the heart observed in IC stroke animals were prevented by rilmenidine treatment. This effect is consistent with previous findings in rats and with rilmenidine is mechanism of action, whose primary site of action is the I1 receptors in the RVLM. Activation of these receptors inhibits central sympathetic outflow and, consequently, prevents the sympathetic hyperactivity induced by IC stroke in the treated group.

Regarding the QTc interval, the ECG analysis showed a significant increase in stroke animals. This prolongation is a known risk factor for arrhythmias and can be associated with conditions like Takotsubo syndrome or neurogenic myocarditis; however, rilmenidine treatment did not reverse this alteration. Meanwhile, QTc interval in the treated group was not statistically different from that of the control group.

The significant and selective increase in norepinephrine (NE) in cardiac tissue is an important marker of cardiac sympathetic hyperactivity. This finding corroborates that IC stroke disrupted central autonomic control, leading to increased peripheral NE release. NE is the primary catecholamine released by postganglionic sympathetic nerve terminals directly in the myocardium, and its elevation in cardiac tissue reflects increased sympathetic nervous activity directed to the heart (Wong, 2006; Xu and Li, 2015; Goldstein, 2025).

Sympathetic hyperactivation acts as a potent pro-inflammatory signal that contributes to the transmigration of immune cells (Kopp, 2015). The hearts of untreated stroke animals showed a mild infiltration of CD68^+^ macrophages and Ly6G^+^ neutrophils. The presence of both cell populations, in a context of sympathetic hyperactivity and stroke, may be associated with the onset of neurogenic myocarditis. Excessive NE release is cardiotoxic and can cause cardiomyocyte necrosis, which, in turn, can lead to the recruitment of macrophages and neutrophils (Hu et al., 2023; Roth et al., 2023). The Ly6G^+^ neutrophil infiltration is characteristic of the acute phase of inflammation, while the presence of CD68^+^ macrophages suggests the beginning of the repair phase or the persistence of the inflammatory stimulus; therefore, their presence can indicate a cardiac inflammatory response (Hu et al., 2023). This finding represents an important similarity with what is observed in Takotsubo Syndrome (Broken Heart Syndrome). The Takotsubo Syndrome is characterized by acute ventricular dysfunction triggered by intense emotional or physical stress, mediated by a massive catecholamine discharge, and is associated with inflammatory infiltration in cardiac tissue (Lyon et al., 2008).

The rilmenidine, by acting on I1 receptors present in structures like the RVLM, can reduce stroke-induced sympathetic hyperactivity, preventing excessive NE release to the heart. This action may impede the primary trigger for the expression of substances such as chemokines and adhesion molecules that can activate inflammatory pathways in the myocardium. Furthermore, by normalizing sympathetic tone, rilmenidine may also be acting on the integrity of the cardiac vascular endothelium, making it less permeable and prone to infiltration by immune system cells (Yalçın et al., 2023; Hua et al., 2025).

The insular stroke model established in C57BL/6J mice demonstrates key aspects of face, construct, and predictive validity, supporting its translational relevance (Pankevich et al., 2013). Microinjection of autologous blood into the IC reproduces the pathological features and blood extravasation observed in spontaneous intracerebral hemorrhage. Moreover, the model successfully reproduces key pathophysiological mechanisms, such as the development of sympathetic hyperactivity and its cardiovascular repercussions. In addition, the model exhibits responsiveness to pharmacological intervention, as rilmenidine treatment attenuated the post-stroke alterations, supporting its utility for investigating therapeutic strategies aimed at central sympathetic modulation. In summary, the results obtained demonstrate that the IC stroke model in mice induces significant sympathetic hyperactivation, characterized by tachycardia, increased NE, and infiltration of immune cells in the myocardium. Rilmenidine treatment normalized these parameters, preventing cardiac injury indicating that the toxic sympathoexcitation is centrally mediated and may involve RVLM neuronal overactivity. These findings validate the IC stroke model in mice as a relevant and important strategy for investigating mechanisms associated with insular stroke. Additionally, the study demonstrates the efficacy of rilmenidine in reducing these stroke-induced injuries.

## Ethical standards

Experiments have been approved by the CEUA-UFMG (protocol 233/2023) ethics committee and performed in accordance with the U.S. NIH Guide for the Care and Use of Laboratory Animals.

## Acknowledgements / Sources of funding

We thank Fundação de Amparo à Pesquisa do Estado de Minas Gerais (FAPEMIG APQ- 01128-21; APQ-01154-23; APQ-05838-23, #RED0008123, APQ-04203-23), Conselho

Nacional de Desenvolvimento Científico e Tecnológico do Brasil (CNPq MAPF PQ 301556/2026-1; ACVG 165774/2021-5; CNPq INCT 406792/2022-4, CNPq

#442473/2023-0, CNPq/MCTI/FNDCT #444251/2024-3), and Serrapilheira Institute (R- 2211-42259). We also acknowledge Nícia Pedreira Soares for technical assistance.

## Data availability

The data that support the fundings of the study is available from the corresponding author upon reasonable request.

